# MarpolBase: Genome database for *Marchantia polymorpha* featuring high quality reference genome sequences

**DOI:** 10.1101/2025.03.30.646155

**Authors:** Yasuhiro Tanizawa, Takako Mochizuki, Masaru Yagura, Mika Sakamoto, Takatomo Fujisawa, Shogo Kawamura, Eita Shimokawa, Shohei Yamaoka, Ryuichi Nishihama, John L. Bowman, Frederic Berger, Katsuyuki Yamato, Takayuki Kohchi, Yasukazu Nakamura

## Abstract

The liverwort *Marchantia polymorpha* is a key model organism for understanding land plant evolution, development, and gene regulation. To support the growing demand for high-quality genomic resources, we present MarpolBase, a comprehensive and integrated genome database that hosts newly assembled, high-accuracy reference genomes for both the male Tak-1 and female Tak-2 accessions, designated as ver. 7.1 reference genomes. These new assemblies, generated using PacBio HiFi long-read sequencing, represent nearly telomere-to-telomere chromosome-level genomes, with improvements in assembly continuity, annotation accuracy, and structural resolution—especially for repeat-rich regions and sex chromosomes.

MarpolBase offers not only access to genome sequences and gene annotations but also provides a unified platform for data exploration, comparative analysis, and community-driven gene nomenclature. It includes keyword-searchable gene pages with structural and functional annotations, expression data integration, genome browser visualization, and online analytical and utility tools. By unifying genome assembly, annotation, nomenclature, and analysis tools in a single platform, MarpolBase serves as a central resource for functional genomics and evolutionary studies in *M. polymorpha*, and a model for future plant genome databases.

## Introduction

The liverwort *Marchantia polymorpha* is increasingly recognized as a pivotal model organism for studying plant evolution, development, and genomics. As a plant species in the bryophyte lineage, sister to the vascular plant (tracheophyte) lineage, *M. polymorpha* can provide crucial insights into the adaptations that facilitated the evolution of land plants from an algal ancestor (Kohchi *et al*. 2021; Bowman *et al*. 2022). Unlike many vascular plants, liverworts are believed to have not undergone ancient whole-genome duplication (WGD) events, resulting in low genetic redundancy across most regulatory pathways (Bowman *et al*. 2017), which facilitates clearer functional analyses of regulatory networks and gene functions. In addition to its simple body plan, *M. polymorpha* exhibits a dominant haploid gametophytic phase alternating with a short diploid sporophytic phase, enabling direct observation of gene function without the complications of heterozygosity. To take advantage of these features, various molecular genetic tools have been developed over the past decades, including *Agrobacterium*-mediated transformation (e.g., the AgarTrap method) (Ishizaki *et al*. 2008; Tsuboyama *et al*. 2018), CRISPR/Cas9 genome editing (Sugano and Nishihama 2018; Sugano *et al*. 2018), microRNA-based gene knockdown strategies (Flores-Sandoval *et al*. 2016), optimized technique for constitutive transgene expression (Tse *et al*. 2024), and a conditional gene expression/deletion system using an endogenous heat-shock promoter and Cre/*lox*P site-specific recombination (Nishihama *et al*. 2016). This extensive molecular tool set has made *M. polymorpha* a powerful system for evolutionary, developmental, and functional genomics studies.

In the *Marchantia* research community, the male Takaragaike-1 (Tak-1) and female Takaragaike-2 (Tak-2) accessions, both belonging to *M. polymorpha* subsp. *ruderalis*, have been widely used as reference laboratory strains, and their genomes have been sequenced to establish reference assemblies. The genome of *M. polymorpha* consists of eight autosomes and a single sex chromosome (U in females and V in males). Previous estimates suggested a genome size of approximately 220 Mb, containing around 20,000 predicted protein-coding genes (Bowman *et al*. 2017; Montgomery *et al*. 2020). The first genome sequencing project for *M. polymorpha* began in 2008 as part of a U.S. Department of Energy’s Joint Genome Institute (JGI) initiative. The whole-genome sequence was determined using a BC4 accession, in which the female Tak-2 accession had been backcrossed to the male Tak-1 accession four times. The initial draft genome, ver. 0.6, was released to the research community in 2011, followed by the first publicly available reference genome, ver. 3.1 (JGI3.1), which consisted of approximately 3,000 scaffold sequences (Bowman *et al*. 2017).

The next major milestone was achieved in 2020 (MpTak1_v5.1), which integrated long-read sequencing technology and Hi-C scaffolding, enabling chromosome-scale genome assembly (Montgomery *et al*. 2020). Subsequently, in 2021, the Tak-2 sex chromosome (chrU) sequence was determined (Iwasaki *et al*. 2021) and, together with the Tak-1 nuclear genome sequences, published as the standard reference genome, ver. 6.1 (MpTak_v6.1), which was designed for analyses that do not require consideration of sex differences. Further refinement of gene models and annotations led to the release of MpTak_v6.1r2 in 2023, which remained the most up-to-date genome until the release of ver. 7.1 described in this paper. In addition to these reference genomes, assemblies from other subspecies have also been published, including MppBR5 (*M. polymorpha* subsp. *polymorpha* strain BR5) and MpmSA2 (*M. polymorpha* subsp. *montivagans* strain SA2) (Linde *et al*. 2020), further expanding genomic resources for *M. polymorpha* research.

The *Marchantia* Genome Database, MarpolBase, was originally developed as a support Wiki site to aggregate information for the ver. 3.1 genome project but later evolved into a fully functional genome database following the release of ver. 5.1. It provides a suite of tools for genome browsing, gene annotation, and expression data visualization, fostering data sharing and collaborative research within the research community. MarpolBase is also seamlessly integrated with the *Marchantia* Expression Database (MBEX) (Kawamura *et al*. 2022) through embedded visualizations in gene detail pages, which has been widely used to explore transcriptomic data across various developmental stages and experimental conditions (Minamino *et al*. 2023; Yamamoto-Negi *et al*. 2024). As a key feature, MarpolBase also hosts the *Marchantia* Nomenclature Database, which allows researchers to register gene names to promote consistency and reduce both redundancy and confusion in scientific communication. To ensure standardization and avoid conflicts, we strongly recommend registering gene names in accordance with the naming guideline (Bowman *et al*. 2016) before publishing research findings.

In this paper, we present a high-quality genome sequencing effort using PacBio HiFi long-read technology, generating nearly telomere-to-telomere assemblies for both Tak-1 and Tak-2, collectively referred to as the ver. 7.1 genomes. The new genome assemblies significantly enhance both completeness and accuracy by resolving previous gaps and errors. Additionally, this paper describes the development of an updated version of MarpolBase, which integrates the ver. 7.1 genome and associated genomic resources. These advancements provide a refined genomic foundation for future research in plant evolutionary biology, developmental genetics, and functional genomics, further solidifying *M. polymorpha* as a premier model system for land plant studies.

## Results and discussion

### Genome sequencing and assembly

The genome assembly as well as gene annotation was performed following the workflow shown in Fig. 1. PacBio HiFi whole-genome sequencing of the male Tak-1 and female Tak-2 accessions generated a total of 38.5 Gb for Tak-1 and 37.2 Gb for Tak-2, with an average read length of 14 kb and a maximum read length of 51 kb. These sequencing data correspond to an approximate depth of coverage of 160× for each accession. After filtering out reads derived from organellar genomes (chloroplast and mitochondrial DNA), the datasets for Tak-1 and Tak-2 had effective coverages of approximately 106× and 108×, respectively, and were subsequently used for genome assembly (Supplementary Table S1). The k-mer (k = 21) distribution plots exhibited a distinct single peak at 100× genome coverage, indicating the presence of high-quality reads derived from the haploid genome. The estimated genome sizes were 217.8 Mb for Tak-1 and 221.3 Mb for Tak-2, both slightly smaller than the actual assembled genome sizes (∼240 Mb) (Supplementary Fig. S1).

**Fig. 1.**
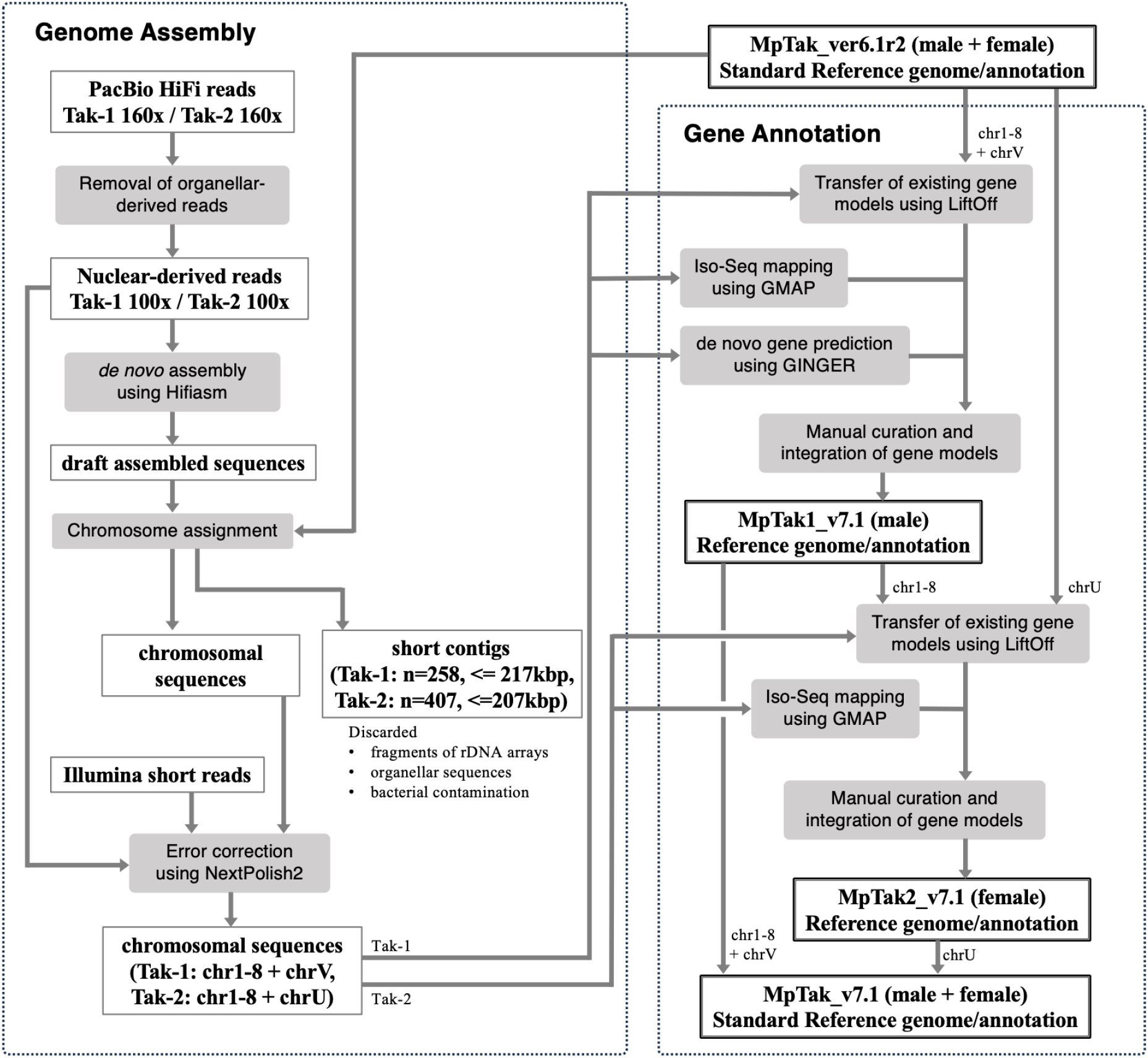
Workflow diagram of genome assembly and gene annotation

Genome assembly was performed using Hifiasm v0.19.5, followed by chromosome assignment through the sequence alignment against the ver. 6.1 reference genome. Specifically, chromosome 3 of Tak-1 and chromosome 4 of Tak-2 were each split into two non-overlapping contigs, which were subsequently joined by introducing a gap. Short sequences that could not be assigned to any chromosome were identified as fragments of organellar genomes, rDNA regions, or bacterial contaminants and were therefore removed. To further improve sequence accuracy, the assembled genomes were polished using NextPolish2 (v0.1.1) with Illumina short reads and PacBio HiFi reads, following the recommended procedure of the software. As a result, nine chromosome-level pseudomolecules were obtained for both Tak-1 and Tak-2, which were designated as ver. 7.1 (MpTak1_v7.1 and MpTak2_v7.1). The final assembly sizes were around 240 Mb, which were approximately 10% larger than the estimated genome sizes as well as the previous versions of the reference genomes. The discrepancy in genome size between the estimated value and that of the previous assembly may arise from the underrepresentation of high-copy repeats in K-mer–based analysis and the potential collapse of repetitive regions during genome assembly.

Gene annotation was primarily lifted over from ver. 6.1 to ver. 7.1 using the Liftoff tool (v1.6.3) followed by manual curation, which will be described in detail later. Genome statistics and comparisons with the previous version are summarized in Table 1.

**Table 1.** Statistics of the *M. polymorpha* ver. 7.1 reference genomes and annotation as well as comparison with the ver. 6.1 genome.

|  | MpTak1_v7.1 | MpTak2_v7.1 | MpTak_v7.1 | MpTak_v6.1 |
| --- | --- | --- | --- | --- |
| **Genome sequence** |  |  |  |  |
| Number of sequences | 9<br>(chr1-8, chrV) | 9<br>(chr1-8, chrU) | 10<br>(chr1-8, chrU, chrV) | 39<br>(chr1-8, chrU, chrV,<br>29 unplaced contigs) |
| Total length | 242,451,485 bp | 241,484,302 bp | 248,042,680 bp | 220846659 bp |
| # of gaps / total gap length | 1 / 500 bp | 1 / 500 bp | 1 / 500 bp | 1,055 / 521,279 bp |
| Repeat ratio | 35.8% | 35.8% | 36.8% | 30.9% |
| BUSCO completeness (genome) | 99.6% | 99.6% | 99.6% | 99.6% |
| K-mer completeness | 99.6% | 99.6% | - | 93.9% * |
| Quality value | 82.9 | 82.9 | 99.6% | 44.7 * |
| **Gene annotation** |  |  |  |  |
| Number of coding loci | 17,944 | 18,136 | 18,007 | 18,038 |
| Number of coding sequence<br>(incl. alternative transcripts) | 22,172 | 22,321 | 22,248 | 22,674 |
| BUSCO completeness (proteome) | 99.6% | 99.6% | 99.6% | 99.6% |

\* Calculated against the genome assembly excluding chrU using Tak-1 Illumina reads

BUSCO analysis using single-copy orthologs (BUSCO v5.8.2 with the eukaryota_odb10 dataset) confirmed that completeness was already high in ver. 6.1, with no further improvements observed in ver. 7.1. However, k-mer completeness and QV scores showed significant improvements, suggesting a more accurate reconstruction of intergenic regions, including repetitive elements. The comparison with previous versions revealed no major structural differences in autosomal sequences, and the Hi-C contact map also confirmed the consistency of the chromosome structure of the ver. 7.1 genomes (Supplementary Fig. S2 and S3). These results highlighted that the previous reference genome (MpTak_v6.1) was already of high quality regarding gene content and chromosome structure.

### Genomic features of ver. 7.1 reference genomes

All assembled chromosome sequences contained either telomeric tandem repeat motifs (TTTAGGG) or rDNA regions at both ends. It is known that nucleolar organizer regions (NORs), containing arrays of rDNA repeats, often form in the region close to telomeres (McStay 2016). Therefore, while the ver. 7.1 assemblies are not telomere-to-telomere in the strictest sense, a nearly complete genome assembly was achieved for both the male and female accessions, with only a single gap remaining in each genome. Centromeres were predicted using CentroMiner, and the results were validated by BLASTN searches using previously identified centromeric repeat sequences as a query sequence (Montgomery *et al*. 2020). With the exception of the male sex chromosome, the centromeres were located in the central regions of each chromosome. Although minor differences in centromere length were observed between the Tak-1 and Tak-2 genomes, their positions were largely conserved, suggesting structural stability of centromeres across sexes. Further analysis of the enrichment of the centromeric histone CENH3 (Takeuchi *et al*. 2024) would be desirable to determine the precise positions of the centromeres. A Circos plot summarizing chromosomal features is shown in Fig. 2, and detailed genomic features, including centromere coordinates and telomere annotations, are listed in Supplementary Table S2.

**Fig. 2.**
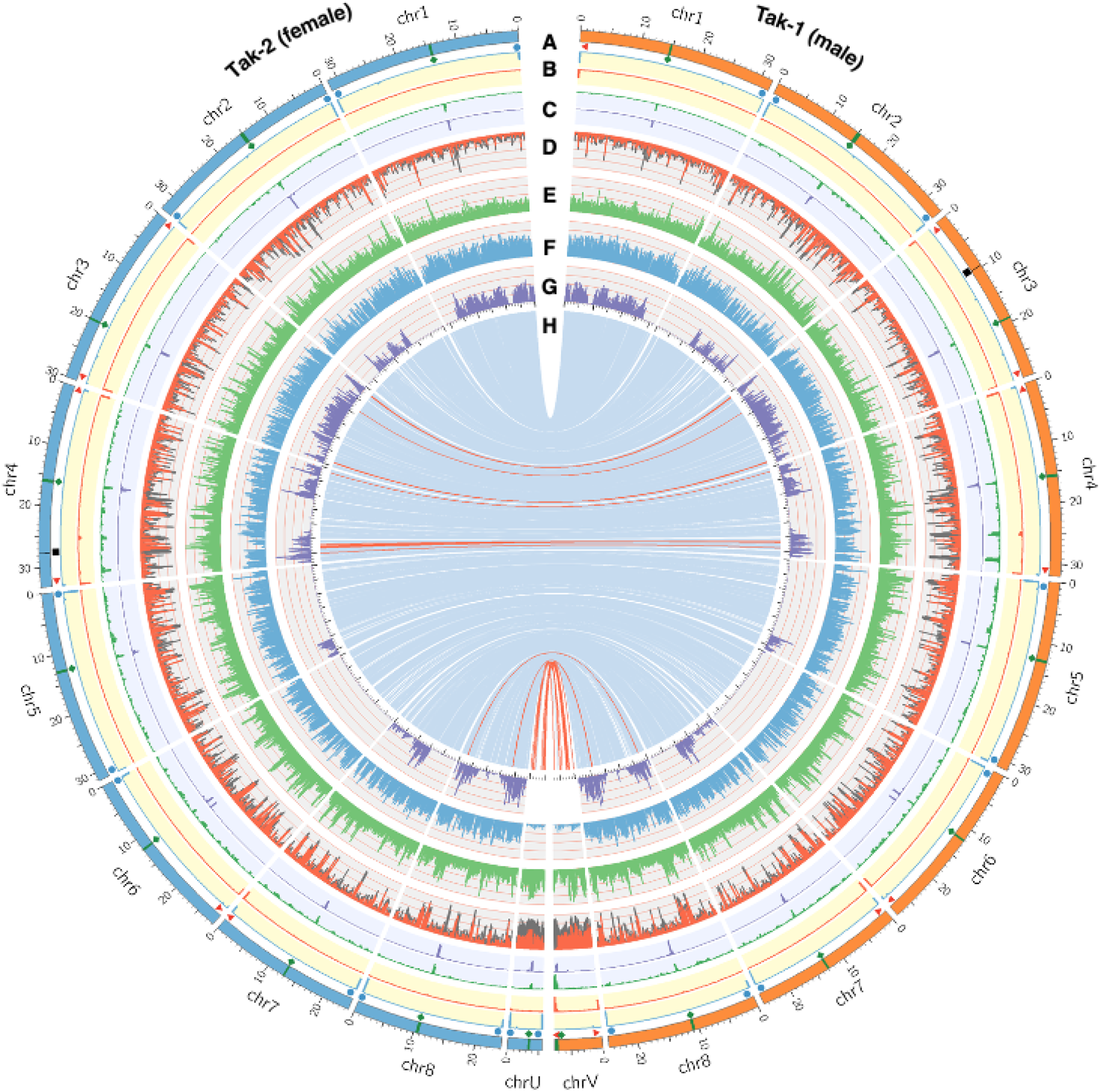
Genomic features of *Marchantia polymorpha* Tak-1 and Tak-2. **A**) Chromosomal features: telomere (red triangle), centromere (green diamond), rDNA cluster (blue circle), assembly gap (black square). **B**) Distribution of rDNAs (blue) and telomeric repeats (red) **C**) Distribution of centromere repeats identified by CentroMiner (green) and BLAST search (purple). **D**) Distribution of repeats, LTR (red) and others (grey) **E**) GC content (y-axis range: 35-55%). **F**) Number of protein coding loci per 200 kbp bin. **G**) Number of variants between Tak-1 and Tak-2 per 200 kbp bin. **H**) Single-copy orthologous gene pairs between Tak-1 and Tak-2, with inverted pairs in red.

A total of 17,423 orthologous gene pairs were identified between Tak-1 and Tak-2 on the autosomes, forming clear 1:1 relationships. No large-scale structural differences were observed, and overall synteny was well-conserved in autosomes between Tak-1 and Tak-2, although small local inversions and insertions/deletions were detected in several regions (Fig. 2H, red lines; Supplementary Fig. S4). In contrast, no clear syntenic relationship was observed between the sex chromosomes (chrV and chrU), and only 21 orthologous gene pairs were detected, many of which correspond to previously reported gametologs (Iwasaki *et al*. 2021).

The only assembly gap in the Tak-1 genome was located at 9.72 Mb on chromosome 3. This region contains a tandem duplication of a gene encoding glutamine synthetase (Mp3g09300). The repeat unit extended for at least 60 kb, and despite the average genome-wide coverage being approximately 100×, this region showed more than 5-fold higher read depth, suggesting that the actual repeat length is substantially greater (Supplementary Fig. S5). Due to the unresolved copy number of the duplicated genes, provisional gene IDs were assigned (e.g., Mp3g09300_L1, see the later section for details). Notably, no tandem duplication was observed at the corresponding region in the Tak-2 genome. Similarly, a large assembly gap on chromosome 4 of the Tak-2 genome at approximately 27.7 Mb corresponded to a highly repetitive region spanning at least 200 kb, which was absent from the Tak-1 genome. This region includes several tandemly duplicated genes, annotated as peroxidases and ion transporters, though their functions remain unknown (Supplementary Fig. S6).

Importantly, the ver. 7.1 assemblies revealed regions on the sex chromosomes that likely reflect mis-assemblies in the previous reference genomes (Supplementary Fig. S7). In our earlier study reporting the chromosome-scale genome assembly (ver. 5.1), we noted the presence of rDNA sequences on the male sex chromosome (chrV) (Montgomery *et al*. 2020). However, no rDNA was identified on chrV in ver. 7.1, which is consistent with the result in earlier cytological analyses of the male sex chromosome (Yamato *et al*. 2007). Regarding the female sex chromosome (chrU), we identified a previously uncharacterized triplicated repeat structure in the middle of the chromosome, which was not observed in ver. This triplication was also reconstructed when using an alternative assembler, Canu, and the uniform read coverage across the region supported its accurate assembly. However, the presence of such repeats may cause mapping ambiguity for short-read sequencing, potentially complicating downstream analyses in this region.

### Improvement of gene annotation

The gene annotation of the *M. polymorpha* reference genome has been inherited from the original annotation of the ver. 3.1 genome, and subsequently maintained through successive versions. During this period, several rounds of updates, including manual curation efforts, were carried out to improve annotation accuracy.

The most extensive update took place during the Genome Annotation Jamboree held as part of the *Marchantia* Workshop in Sendai, Japan, in 2019, where 70 researchers participated in improving gene annotations. As a result, a total of 4,109 gene models were manually curated, including modifications and additions to existing models, as well as the removal of spurious gene models. These improvements were incorporated into an updated version of the genome, MpTak1_v5.1r2, released in 2021.

For the ver. 7.1 genomes, annotation was lifted-over from MpTak_v6.1r2 using the Liftoff tool, which itself was based on MpTak1_v5.1r2 with additional small-scale curation. In Tak-1, 2,011 gene models were manually curated, including 784 deletions and 392 new gene predictions, the latter primarily based on *de novo* gene prediction and full-length transcript mapping. Consequently, the number of annotated genes has slightly decreased in ver. 7.1 compared to ver. 6.1r2. Annotation for Tak-2 was lifted-over from the curated Tak-1 gene models using Liftoff, and an additional 325 genes were supplemented. Most of them were extra copies of existing genes, but 15 genes were newly annotated on the triplicated repeat region of the U chromosome by manual curation, although their functions remain unknown. The lists of genes newly added or removed in Tak-1 and Tak-2 are provided in Supplementary Tables S3 and S4, respectively. Functional assignments to annotated gene models were carried out by searching against multiple reference databases, including KEGG, KOG, and InterPro. These annotations are searchable via the keyword search function in MarpolBase.

Notably, in ver. 7.1, all previously unplaced genes (n = 39), which had resided on scaffolds not assigned to chromosomes (unplaced scaffolds) in earlier versions, were successfully anchored to specific chromosomes. Gene identifiers were reassigned according to the chromosome to which each gene was mapped. One such example is Mp*LRL*, a gene known to play key roles in many aspects of plant growth and development, including rhizoid and reproductive development (Breuninger *et al*. 2016; Saito *et al*. 2023). Until ver. 5.1, the Mp*LRL* locus (Mpzg01410) was fragmented on an unplaced scaffold. Although the entire sequence was recovered in ver. 6.1, it remained on an unplaced scaffold. In ver. 7.1, it was mapped to the 5′ region of Mp2g20700 on chromosome 2 and reassigned the new ID Mp2g20695 (Supplementary Fig. S8). In rare cases, genes that were transferred to a different chromosome were assigned new identifiers to reflect their new chromosomal locations (Supplementary Table S3). For regions affected by local inversions, the gene order in ver. 7.1 is reversed relative to ver. 6.1, but gene identifiers have been preserved for consistency.

### Gene identifier system

The current gene identifier system was introduced with the release of the chromosome-scale genome assembly (ver. 5.1) and is referred to as the MpGene ID system. In this system, each gene locus is assigned with a unique, stable identifier, modeled after the AGI (*Arabidopsis* Genome Initiative) locus code system used in *Arabidopsis thaliana* (Berardini *et al*. 2015). An MpGene ID consists of the prefix “Mp”, followed by the chromosome number (1–8, U, or V), the letter “g” indicating a gene, and a five-digit number (e.g., Mp3g09300).

These five-digit numbers were originally assigned in the order in which genes appear along each chromosome. However, due to local inversions or updates in genome assemblies, gene number order may no longer strictly reflect physical positions in the later versions including ver. 7.1. To distinguish alternative transcripts originating from the same locus, an additional suffix is appended to the gene ID, separated by a period (.), and referred to as the transcript ID (e.g., Mp3g09300.1, Mp3g09300.2).

MpGene IDs are designed as permanent identifiers and are retained across genome versions unless changes are justified by exceptional circumstances. However, the isoform number appended to the gene ID (e.g., .1, .2) is not preserved across versions. This is because transcript isoforms have frequently been added, removed, or revised during annotation updates, making it difficult to maintain consistent isoform numbering. Of note, a subset of genes was assigned provisional IDs in ver. 7.1, which can be identified by the inclusion of an underscore (_) in the gene ID. These provisional IDs fall into two categories:

1. Genes in the assembly gap region of chromosome 3 in the Tak-1 genome, which form tandem repeats and whose precise copy number remains unresolved (e.g., Mp3g09300_L1, Mp3g09295_R2; total of 9 genes).
2. Extra gene copies that appeared during the lift-over from the Tak-1 to the Tak-2 genome, for which no clear orthologous loci exist in Tak-1 (e.g., Mp1g90010_P, Mp1g90020_P; total of 310 genes).

In earlier versions (ver. 5.1 and ver. 6.1), genes located on unplaced scaffolds were given identifiers prefixed with “Mpzg” (e.g., Mpzg01410). However, in ver. 7.1, these genes were successfully assigned to chromosomes and the Mpzg system was retired.

In the earlier ver. 3.1 genome, a different identifier system called Mapoly ID (e.g., Mapoly0010s0028) was used. To support continuity and data integration, a Gene ID Converter is available on MarpolBase to convert between Mapoly IDs, MpGene IDs, and versions from ver. 5.1 to ver. 7.1. Additionally, correspondence tables for gene IDs are available for download.

### Database features

MarpolBase is a comprehensive genomic resource for *M. polymorpha*, designed to support genome annotation, comparative genomics, and functional studies. It consists of a main web platform implemented in Python, integrated with a Docker-based genome browser (WebApollo/JBrowse) and BLAST server (SequenceServer) (Lee *et al*. 2013; Priyam *et al*. 2019). Originally developed to host the genome data for the ver. 5.1 genome derived from *M. polymorpha* subs. *ruderalis*, MarpolBase has since expanded to include multiple genome assemblies including those for other subspecies. The full list of available genomes is provided in Table 2, with MpTak_v7.1 designated as the default reference genome, which is used for searches and analyses unless otherwise specified. The screenshots of MarpolBase are shown in Fig. 3.

**Table 2.** List of available genomes in MarpolBase.

| Name | Description | Number of sequences | size | Protein coding loci | Release | INSDC Assembly ID |
| --- | --- | --- | --- | --- | --- | --- |
| **MpTak_v7.1** | T2T-scale Standard genome (Tak-1 + Tak-2 ChrU) | 10 chromosomes | 248 Mbp | 18,007 | 2024 | GCA_039105155.1 |
| **MpTak1_v7.1** | T2T-scale male reference genome (Tak-1) | 9 chromosomes | 241 Mbp | 17,944 | 2024 | GCA_037833805.1 |
| **MpTak2_v7.1** | T2T-scale female reference genome (Tak-2) | 9 chromosomes | 242 Mbp | 18,136 | 2024 | GCA_037833965.1 |
| **MpTak_v6.1** | Chromosome-scale standard genome (Tak-1 + Tak-2 ChrU) | 10 chromosomes + 261 scaffolds | 223 Mbp | 18,335 | 2021 | (GCA_019973755.1) * |
| **MpTak_v6.1r2** | Same as above with gene annotation revised. | 10 chromosomes + 29 scaffolds | 221 Mbp | 18,038 | 2023 | - |
| **MpTak1_v5.1** | Chromosome-scale male reference genome (Tak-1) | 9 chromosomes + 435 scaffolds | 218 Mbp | 19,421 | 2020 | GCA_009936355.2 |
| **MpTak1_v5.1r2** | Same as above with gene annotation revised. | 9 chromosomes + 435 scaffolds | 218 Mbp | 18,288 | 2021 | - |
| **JGI3.1** | Draft level reference genome sequences (Tak-1 BC4) | 2,957 scaffolds | 225 Mbp | 19,287 | 2017 | GCA_003032435.1 |
| **MppBR5** | Draft genome of subsp. *polymorpha* BR5 | 2,710 scaffolds | 225 Mbp | 17,374 | 2020 | GCA_011319315.1 |
| **MpmSA2** | Draft genome of subsp. *montivagans* SA2 | 2,741 scaffolds | 222 Mbp | 18,806 | 2020 | GCA_011319375.1 |

All genomes were derived from *M. polymorpha* subsp. *ruderalis* except MppBR5 and MpmSA2
\* chrU sequence from Tak-2 is deposited as GCA\_019973755.1

**Fig. 3.**
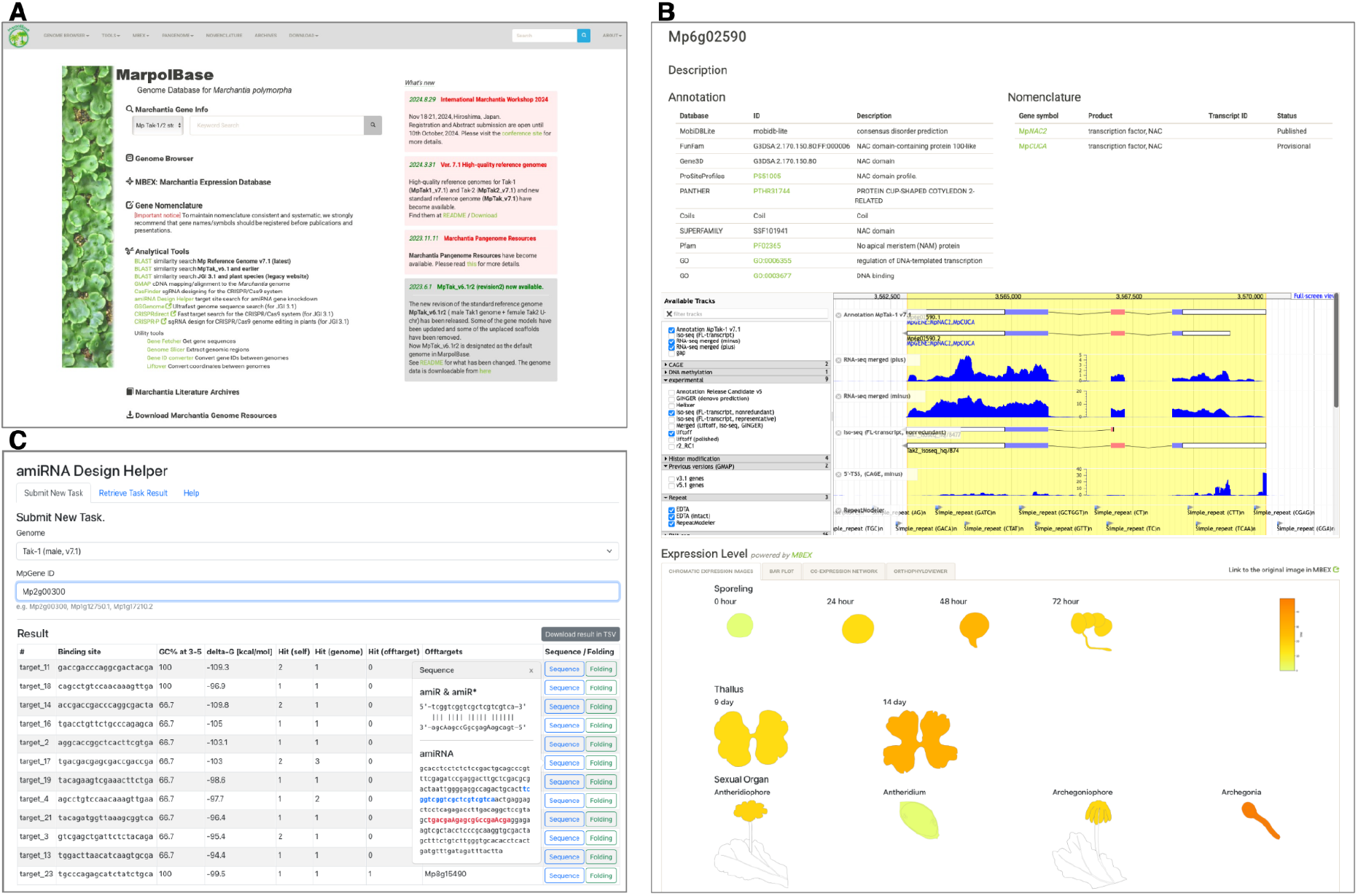
Screenshots of the MarpolBase web interfaces. **A**) Top page of the website. **B**) Gene detail page showing the gene structure in the embedded genome browser and expression pattern retrieved from *Marchantia* Expression Database (MBEX). **C**) amiRNA Design Helper (top: job submission form; bottom: output results), shown as an example of the analytical tools provided by MarpolBase.

The core functionality of MarpolBase lies in its gene search and annotation system, enabling users to retrieve gene-related information through keyword searches and detailed gene pages. These pages provide structural and functional annotations, gene sequences, and visualization tools, along with embedded genome browser views and expression data from MBEX, allowing for interactive exploration of transcriptional profiles. BLAST search results and genome browser links facilitate seamless navigation between different data layers, while integration with the Gene Nomenclature Database ensures consistency in gene naming across studies.

To enhance usability, MarpolBase includes a suite of online analysis tools for sequence comparison, genome annotation, and molecular biology applications. BLAST search enables homology-based sequence retrieval, while GMAP allows splicing-aware alignment of cDNA sequences to the genome. For functional genomics, MarpolBase provides a CRISPR/Cas9 guide RNA design tool, implemented using CASFinder (Aach, Mali and Church 2014), and an artificial microRNA (amiRNA) design tool based on the protocol of Flores-Sandoval *et al*. (Flores-Sandoval *et al*. 2016)), facilitating gene editing and knockdown studies. In addition, utilities for sequence retrieval and gene ID or coordinate conversion between reference genomes are available, ensuring compatibility across different datasets (Table 3).

**Table 3.** List of analytical and utility tools in MarpolBase.

| Tool Name | Description |
| --- | --- |
| **Analytical tools** |  |
| BLAST | BLAST search against the *Marchantia* reference genomes as well as other plant species. Deployed with SequenceServer. |
| GMAP | Splicing-aware alignment of cDNA sequences to the *Marchantia* genome using GMAP. |
| CasFinder | Design of CRISPR/Cas9 guide RNAs based on the CasFinder algorithm. |
| amiRNA Design Helper | Design of artificial microRNA (amiRNA) target sites for gene knockdown studies. |
| **Utility tools** |  |
| Gene Fetcher | Retrieve transcript, CDS, or protein sequences of specific genes. |
| Genome Slicer | Extract custom genomic regions based on chromosomal coordinates. |
| Gene ID converter | Convert gene identifiers between different genome versions. A correspondence table is also available for download. |
| Liftover | Convert genomic coordinates between different genome assemblies. Implemented with transanno. Chain files for local use are also available. |

Beyond genome annotation and analysis, MarpolBase serves as a repository for research data, hosting over 130 accessions of pangenome data (Beaulieu *et al*. 2025). A dedicated genome browser and BLAST search are available for visualizing genetic variants, supporting population and evolutionary studies. Additionally, MarpolBase provides a data-sharing platform upon request, allowing researchers to store and distribute their datasets within the *Marchantia* research community.

### Gene nomenclature database

The *Marchantia* Nomenclature Database, a key component of MarpolBase, provides researchers with a platform to register gene symbols and associated literature for specific loci (Fig. 4). Submitted entries undergo administrator review prior to approval for public access and are linked to gene detail pages. A Naming Guideline for gene nomenclature was published in 2016 (Bowman *et al*. 2016), according to which gene names with the “Mp” prefix are recommended for genes characterized in *M. polymorpha* subsp. *ruderalis*, from which the Tak-1 and Tak-2 reference genomes are derived. Users can select one of two publication statuses: Reserved, where only the gene symbol and minimal information are displayed while linked gene details remain unpublished, or Published, which makes all registered information, including the submitter’s name and literature references, publicly accessible.

**Fig. 4.**
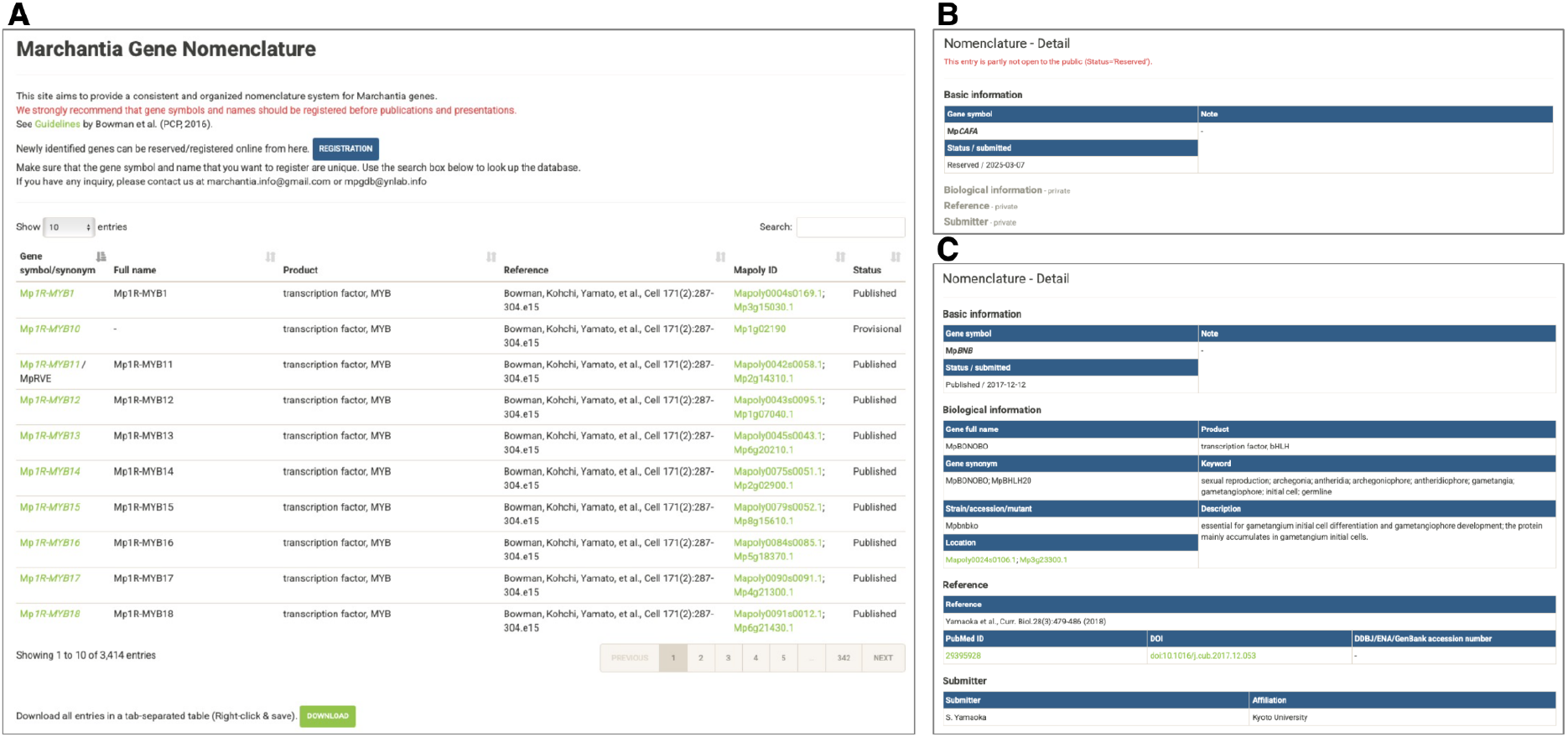
Screenshots of the Gene Nomenclature Database. **A**) Top page showing registered gene names. New gene names can be submitted via the “Registration” button. **B**) Gene detail page with status set to ‘Reserved’. Only minimal information is shown. **C**) Gene detail page with status set to ‘Published’. All information is publicly available.

A lack of standardized nomenclature can lead to confusion, as demonstrated by the case of gene names for RESPIRATORY BURST OXIDASE HOMOLOGs (RBOHs). RBOH proteins are known to be involved in reactive oxygen species production, and *M. polymorpha* possesses two paralogous genes, Mp3g20340 and Mp7g00270 (Kimura *et al*. 2020; Marchetti *et al*. 2021). In 2023, three independent studies on these genes were published almost simultaneously. However, two of these studies assigned the names Mp*RBOH1* and Mp*RBOH2* to the paralogs but in opposite orders, leading to inconsistency (Chu *et al*. 2023; Yotsui *et al*. 2023). Meanwhile, another study followed the nomenclature used in *Arabidopsis* and designated them Mp*RBOHA* and Mp*RBOHB* (Hashimoto *et al*. 2023). Such discrepancies complicate cross-study comparisons and highlight the importance of a unified nomenclature system. More recently, another nomenclature conflict has arisen, in which two independent research groups assigned different gene names, Mp*PLT* and Mp*ANT*, to the same locus, Mp8g11450 (Fu *et al*. 2024; Liu *et al*. 2024).

By utilizing this system, gene name standardization is ensured, preventing redundant or conflicting annotations across studies. It also facilitates proper attribution and citation by associating researchers’ work with specific gene names. The ‘Reserved’ status enables early registration of gene symbols while maintaining the confidentiality of unpublished data, minimizing the risk of unintentional duplication. Furthermore, integrating nomenclature with gene detail pages enhances data visibility, promoting collaboration and efficient information sharing within the research community. To maintain consistency and avoid conflicts in gene nomenclature, we strongly recommend registering gene names prior to publication or presentation of research findings.

## Conclusion

In this study, we present a highly accurate and nearly complete telomere-to-telomere genome assembly of *M. polymorpha* using PacBio HiFi sequencing, resulting in ver. 7.1 reference genomes for both the male (Tak-1) and female (Tak-2) accessions. These assemblies resolve previous assembly gaps and structural ambiguities, particularly in sex chromosomes, and provide improved continuity and accuracy in gene annotations. The updated annotation, supported by manual curation and integration of transcriptomic data, offers a robust framework for gene-level analyses. Importantly, all previously unplaced genes have now been successfully assigned to chromosomes, and repeat-rich regions such as the tandem duplications and centromere-proximal domains were resolved with unprecedented precision.

Alongside the updated genome, we developed an enhanced version of MarpolBase, a comprehensive genomic resource that integrates genome sequence data, gene annotation, expression profiles, functional predictions, and a unified gene nomenclature system. MarpolBase facilitates efficient data exploration, hypothesis testing, and cross-study comparisons by offering searchable gene information, interactive genome browsers, and online analysis tools, including support for CRISPR/Cas9 and miRNA-based gene manipulation. As *M. polymorpha* continues to gain prominence as a model system for studying early land plant evolution and gene function, we anticipate that the ver. 7.1 genomes and the expanded features of MarpolBase will serve as foundational resources for the plant research community.

## Materials and methods

### DNA extraction and genome sequencing

Genomic DNA was extracted from 2-week-old thalli of male and female *M. polymorpha* subsp. *ruderalis* accessions, Tak-1 and Tak-2, using a CTAB-based method followed by purification using QIAGEN Genomic-tip kit (QIAGEN) to obtain high-molecular-weight DNA suitable for long-read sequencing. For HiFi library preparation, DNA was sheared using a g-TUBE (Covaris), and size selection was performed using 35% AMPure PB beads to remove fragments shorter than 3 kb. The SMRTbell Express Template Prep Kit 3.0 was used for library construction, and sequencing was carried out on a PacBio Sequel II system with Sequel II Binding Kit 3.2 and Sequel II Sequencing Kit 2.0. Raw reads were processed using DeepConsensus v1.2 (Baid *et al*. 2023) to generate high-fidelity (HiFi) reads, which were used for downstream genome assembly and analysis.

### Genome assembly

Prior to genome assembly, raw reads were mapped to reference sequences of organellar genomes (chloroplast and mitochondrial DNA) to remove organelle-derived sequences. The organellar genome sequences were obtained from MarpolBase. As both the chloroplast and mitochondrial genomes are circular, additional versions with adjusted starting coordinates were prepared to improve mapping rate. Also, as the inverted repeat (IR) regions of the chloroplast genome are known to undergo flip-flop recombination, an alternative version of the chloroplast genome with the IR-flipped region was also included. Reads were mapped using Minimap2 v0.2-r123 (Li 2018), and reads with ≥90% alignment coverage to the organellar genome were excluded from the dataset.

Genome assembly was performed using Hifiasm v0.19.5 (Cheng *et al*. 2021) with the ‘--primary’ option enabled. To assign contigs to chromosomes, contigs ≥1 Mb in length were compared using D-GENIES (Cabanettes and Klopp 2018) with previous versions of the reference genome (MpTak_v6.1r2 for Tak-1 and an unpublished assembly related to Iwasaki *et al*. (2022) for Tak-2 (Iwasaki *et al*. 2021)). During the assembly process, sequences corresponding to chr4 and chr5 in the Tak-1 genome were found to be fragmented across multiple contigs. To resolve this, an additional assembly was generated using Hifiasm with the ‘--primary --nhap1’ option, and the resulting contigs were merged with the original assembly. Short contigs failed to be assigned to chromosomes (258 contigs for Tak-1, max length: 217 kb; 407 contigs for Tak-2, max length: 207 kb) were screened for contamination or non-nuclear origins. BLASTN (Camacho *et al*. 2009) searches were conducted against bacterial genomes, organellar sequences, and rDNA regions to identify contaminant sequences. Bacterial contamination was assessed using prokaryote representative genomes from RefSeq, while rDNA regions were predicted using Barrnap (Seemann). For comparative purposes, an additional de novo assembly was generated using Canu v2.2 (Koren *et al*. 2017), specifying an estimated genome size of 240 Mb with default parameters. To further improve consensus accuracy, the assembled genomes were error-corrected using NextPolish2 v0.1.1 (Hu *et al*. 2024). This process incorporated Illumina short reads and PacBio HiFi reads following the instruction manual, utilizing Meryl (v1.3), Winnowmap (v2.03), and Yak (v0.1) for polishing.

### K-mer Analysis

K-mer analysis was performed using Jellyfish v2.2.10 with a K-mer size of 21 to count the frequency of all 21-mers in the PacBio HiFi reads. The resulting k-mer frequency histograms were analyzed and visualized using GenomeScope v1.0 to estimate genome size, heterozygosity, and the proportion of repetitive elements.

### Prediction of genomic features

Centromeric regions were predicted using CentroMiner via the quarTeT web server (https://www.atcgn.com/quarTeT/home.html), and the results were confirmed by BLASTN (v2.15.0) searches using previously identified centromeric repeat sequences as queries. Ribosomal DNA (rDNA) regions were predicted using Barrnap v0.9 with default settings. De novo repeat discovery was performed using EDTA v2.0.0 and a combination of RepeatModeler and RepeatMasker implemented in the TeTools v1.87 pipeline. Telomeric repeat motifs (CCCTAAA) were detected using the Telomere Identification toolKit (tidk) v0.2.31, and the predictions were validated with the RepeatMasker output to confirm telomeric localization.

### Gene annotation

For the Tak-1 genome, a total of 22,612 transcript models from the autosomes and chromosome V of MpTak_v6.1r2 were transferred to the ver. 7.1 genome using Liftoff v1.6.3 (Shumate and Salzberg 2021)with the ‘-copies’ option to allow detection of extra gene copies. As a result, 22,690 models were successfully lifted-over, including extra copies. Among these, 22,555 models were mapped in a 1:1 correspondence with a valid open reading frame (valid_ORF=True and extra_copy_number=0), while 9 genes failed to map. To identify novel gene loci, full-length transcript sequences (Iso-Seq) reads obtained from BioProject PRJDB8530 were aligned to the ver. 7.1 genome using GMAP v2023.10.10 (Wu *et al*. 2016). Additionally, *de novo* gene prediction was conducted by GINGER v1.0.1 (Taniguchi *et al*. 2023) with RNA-seq reads (SRA accession no. SRR896227) as input evidence. The outputs from Iso-Seq mapping and GINGER were compared with the Liftoff results using GffCompare v0.12.6 (Pertea and Pertea 2020), and 177 and 1,015 transcript models, respectively, were identified as novel gene candidates not represented in the Liftoff annotation. Exting transcript models that were inconsistent with Iso-Seq alignments, as well as models transferred by Liftoff but lacking valid ORFs, were subjected to manual curation. Manual curation was conducted using the WebApollo platform, and the resulting curated annotation was designated as the reference gene annotation for the Tak-1 ver. 7.1 genome.

For the Tak-2 genome, gene annotation was performed by transferring a total of 22,104 transcript models using Liftoff, consisting of the curated Tak-1 autosomal genes and the chrU (female sex chromosome) annotation from MpTak_v6.1r2. A total of 22,347 genes were successfully mapped to the Tak-2 genome, including extra copies. Of these, 21,807 genes were mapped in a 1:1 relationship, while 55 genes failed to map. As with Tak-1, novel candidates identified from Iso-Seq evidence and genes lacking valid ORFs were manually curated and merged into the final Tak-2 genome annotation.

### Evaluation of genome assembly and annotation

The K-mer-based completeness and consensus quality value (QV) were calculated using Merqury v1.3 based on K-mer counts generated with Meryl v1.3 (k = 21). Assembly and annotation completeness were also assessed using BUSCO v5.8.2 with the eukaryota_odb10 dataset.

To assess chromosome-level assembly structure, Hi-C contact matrices were created as follows. Hi-C sequencing reads were obtained from the INSDC Sequence Read Archive (SRA accession no. SRR9974619). The reads were preprocessed with fastp, and mapped to the assembled genome using Chromap v0.2.5 with the options ‘--preset hic --remove-pcr-duplicates’ to generate contact pair files. These were processed using Juicer Tools v1.22 to create contact matrices, which were visualized using Juicebox v1.11.08.

### Database construction

The main website of MarpolBase was implemented using Python v3.7.6 and the Flask web framework, with MySQL v8.3 as the database management system. The genome browser and BLAST web server were deployed using containerized instances of WebApollo and SequenceServer, respectively. The amiRNA Design Helper tool was developed with a React-based frontend and a backend powered by FastAPI and Redis. Additionally, nginx is used to serve specific components of the data download site.

## Data availablity

The genome sequences and related datasets are available from the MarpolBase download site (https://marchantia.info/download/). In addition, the assembled ver. 7.1 reference genomes and raw sequencing reads have been deposited in the INSDC under BioProject accession number PRJDB16711. The genome assemblies are available under the following accession numbers: MpTak1_v7.1 (GCA_037833805.1), MpTak2_v7.1 (GCA_037833965.1), and MpTak_v7.1 (GCA_039105155.1). The corresponding PacBio HiFi reads are available under DRR504758 (Tak-1) and DRR504759 (Tak-2).

## Acknowledgments

This work was supported by JSPS KAKENHI Grant Numbers 16H06279 (PAGS) and 20K15783, and GteX Program Japan Grant Number JPMJGX23B0. We thank Dr. Hideki Nagasaki for his contributions during the early stages of the project. We also thank Dr. Yoshihiro Yoshitake for sample preparation for genome sequencing, Dr. Facundo Romani for his extensive effort in gene nomenclature, Dr. Yuuki Sakai and Dr. Takehiko Kanazawa for their suggestions regarding the amiRNA Design Helper, and Dr. Takeshi Ara for his support in data submission to DDBJ. Lastly, we are grateful to all members of the *Marchantia* research community who participated in the Genome Annotation Jamboree for their valuable contributions to gene model improvement.

## Notes

### Competing Interest Statement

The authors have declared no competing interest.

