## Supplementary Figures for "MarpolBase: Genome database for *Marchantia polymorpha* featuring high quality reference genome sequences"

**Supplementary Table S1.**
Read statistics.

**Supplementary Table S2.**
Chromosome features, length, number of genes, centromere, presence/absence of telomere/rDNA array.

**Supplementary Table S3.**
Gene models modified/added/deleted while updating from ver. 6.1 to ver. 7.1.

**Supplementary Table S4.**
Gene models added/deleted while transferring annotation from Tak-1 to Tak-2 (ver. 7.1).

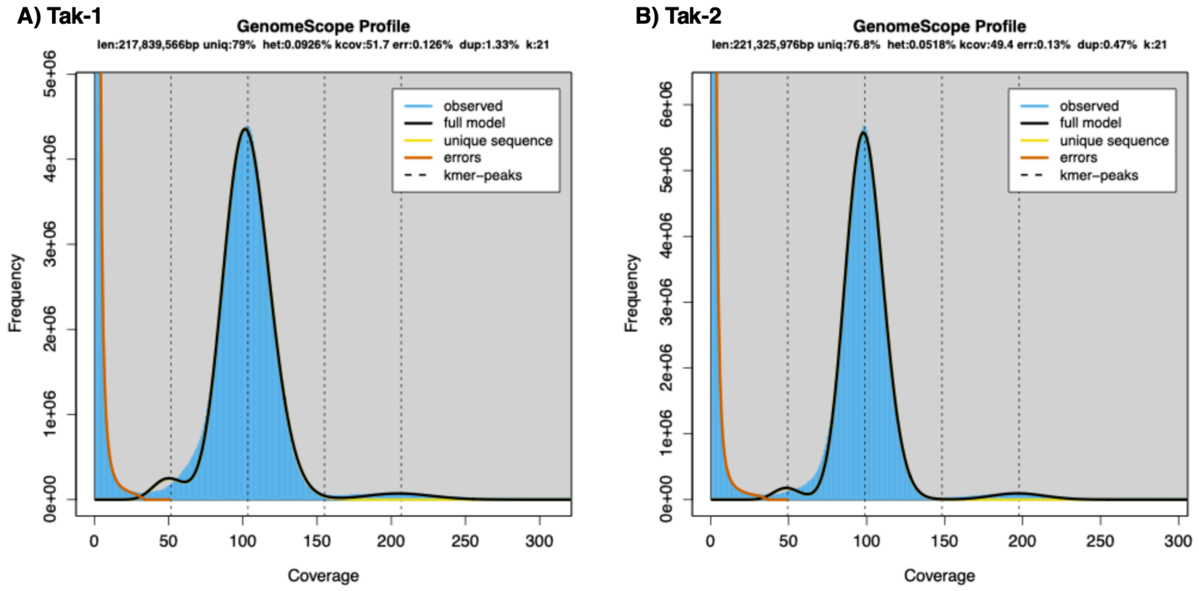


**Supplementary Fig. S1** K-mer distribution and genome size estimation for the Tak-1 (**A**) and Tak-2 (**B**) genomes. K-mer (k = 21) frequencies were calculated using Jellyfish v2.2.10 and analyzed with GenomeScope v1.0 to estimate genome size and repeat content.

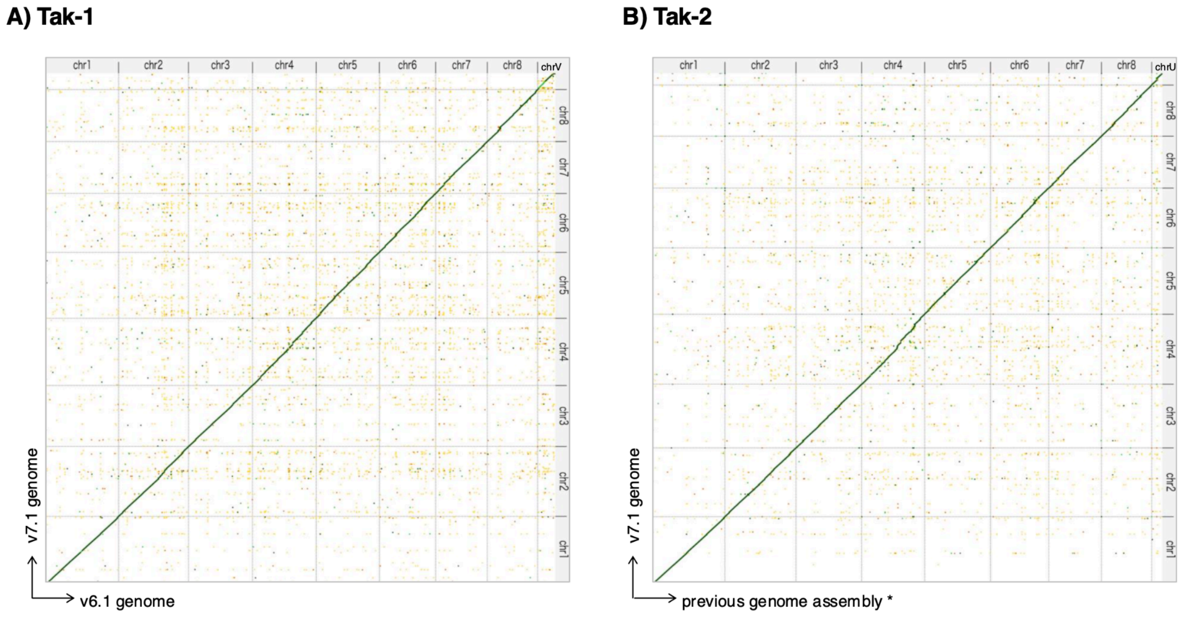


**Supplementary Fig. S2** Comparison with previous versions of the genome assembly: **A**) Tak-1 and **B**) Tak-2. * For the Tak-2 genome, comparison was made with an unpublished assembly related to *Iwasaki et al*. Curr. Biol. (2021), which was constructed using PacBio long reads and Hi-C scaffolding.

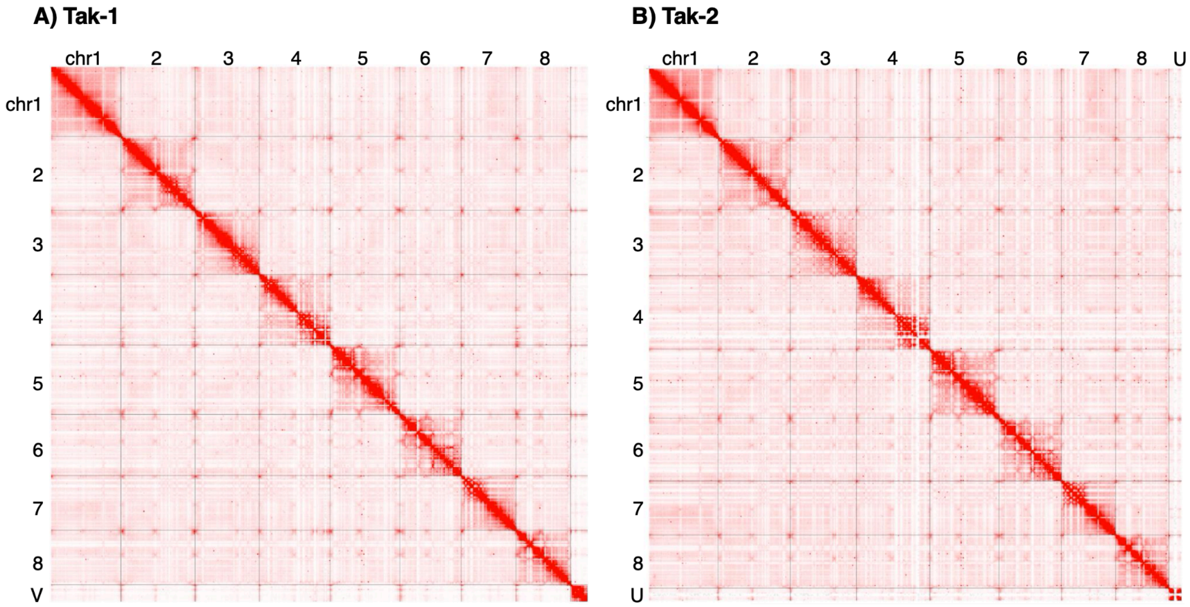


**Supplementary Fig. S3** Hi-C contact maps of the chromosome-level genome assemblies for Tak-1 (**A**) and Tak-2 (**B**). Hi-C reads were mapped to each genome assembly using Chromap and contact matrices were generated using Juicer Tools and visualized with Juicebox. The contact maps support the structural integrity of the assembled chromosomes in both strains.

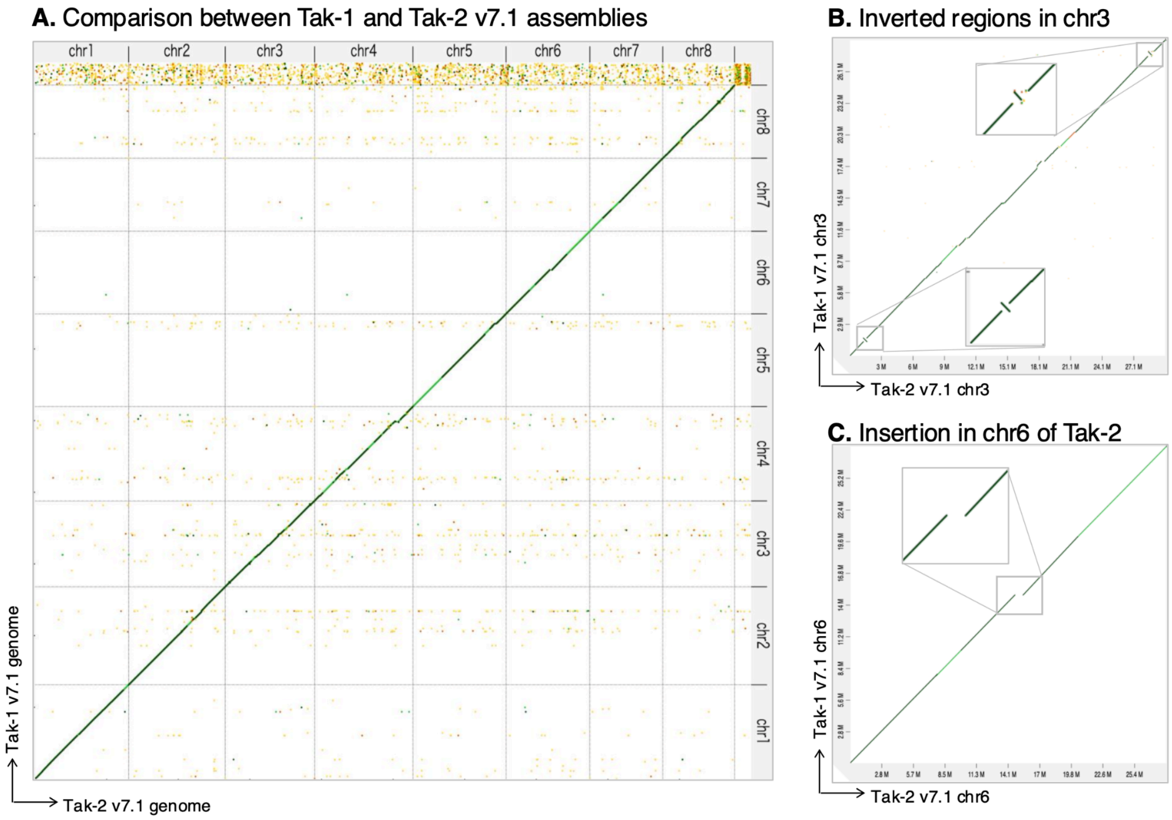


**Supplementary Fig. S4 A**) Comparison between Tak-1 and Tak-2 ver. 7.1 assemblies. No large homologous regions were observed between chrV and chrU. **B**) Examples of local inversions. **1.** chr3 1.31–1.65 Mb in Tak-1 and 1.30–1.64 Mb in Tak-2; **2.** chr3: 27.68–27.96 Mb in Tak-1 and 28.52–28.78 Mb in Tak-2. **C**) Example of an insertion/deletion. **3.** Insertion in chr6 of Tak-2: 14.86 Mb in Tak-1 and 14.73–15.46 Mb in Tak-2.

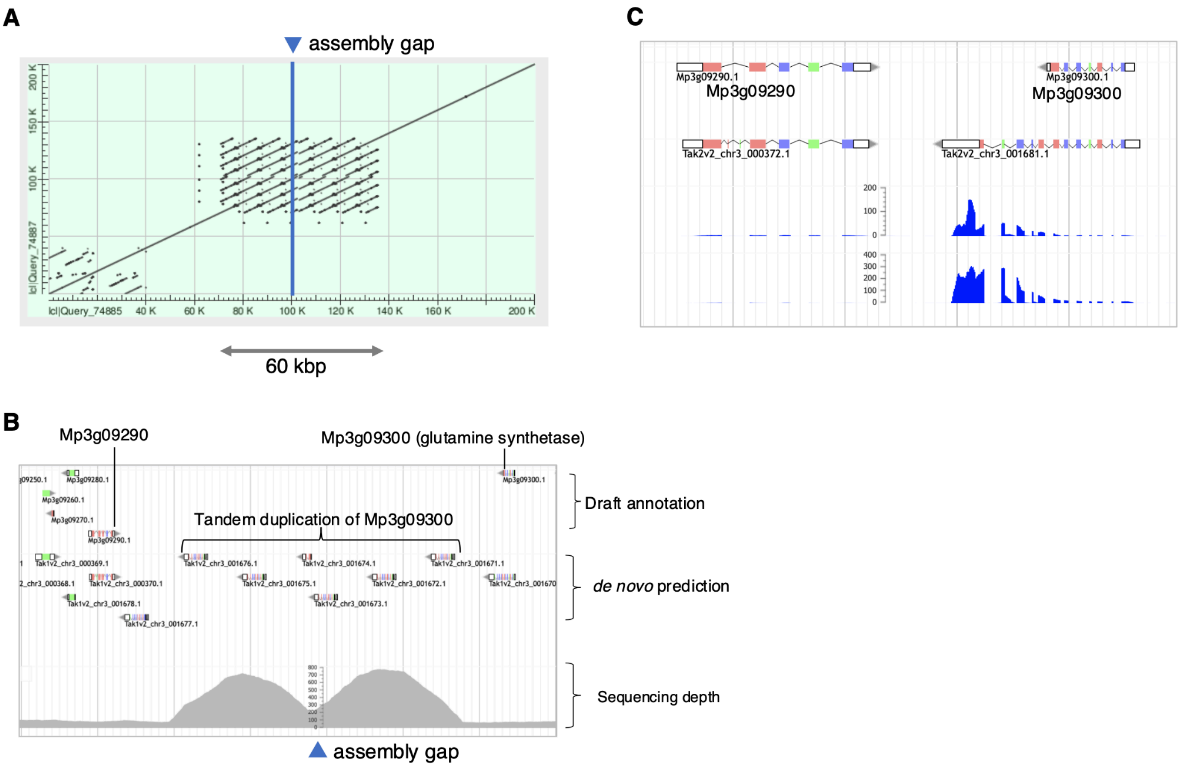


**Supplementary Fig. S5** The region surrounding the assembly gap on chromosome 3 of Tak-1 (at 9.72 Mb). **A**) Self-dot plot visualized using the NCBI BLAST web service, showing a repetitive structure around the gap region. **B**) Gene annotation and sequencing depth in the same region. Sequencing depth is notably higher than in adjacent regions, suggesting a higher copy number of tandem repeats. **C**) The corresponding region on chromosome 3 of Tak-2, which lacks an assembly gap and repetitive structure.

Provisional gene IDs were assigned to the duplicated genes. Genes to the left of the gap: Mp3g09295_L1, Mp3g09300_L1, L2, L3. Genes to the right of the gap: Mp3g09295_R1, R2, Mp3g09300_R1, R2, R3.

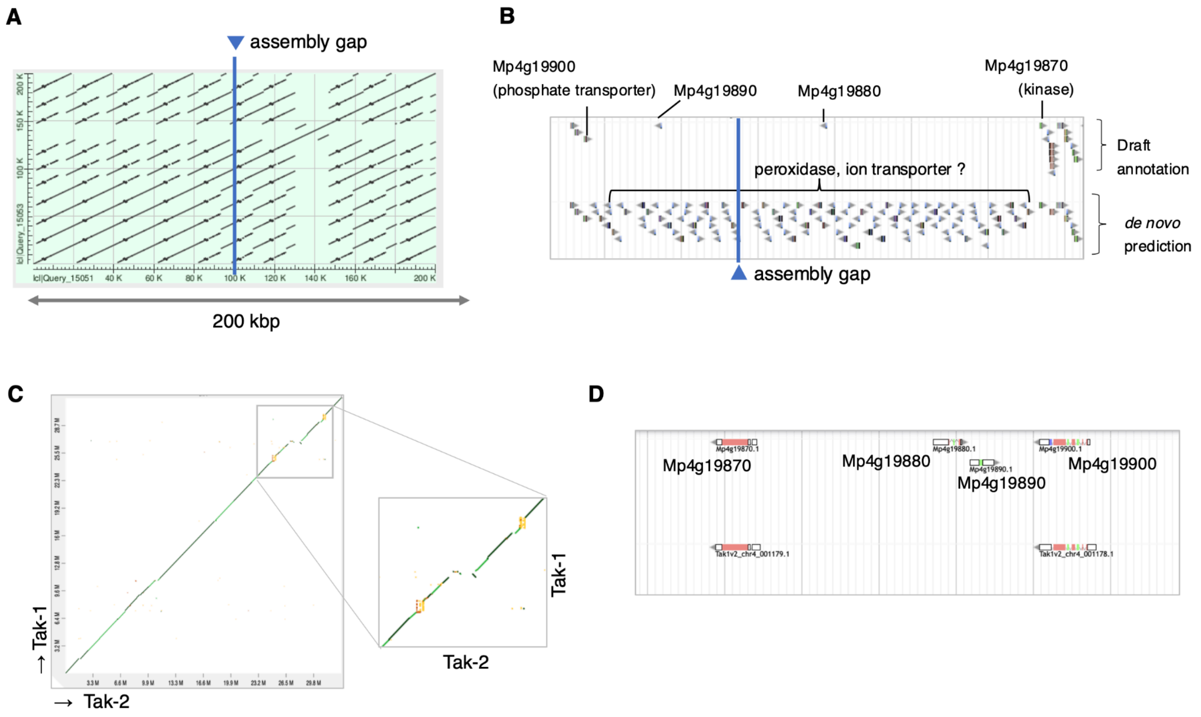


**Supplementary Fig. S6** The region surrounding the assembly gap on chromosome 4 of Tak-2 (at 27.7 Mb). **A**) Self-dot plot visualized using the NCBI BLAST web service, showing a highly repetitive structure near the gap region. **B**) Gene annotation in the same region. Sequencing depth could not be visualized due to excessive read mapping, indicating extremely high repeat content. **C**, **D**) The corresponding region on chromosome 3 of Tak-1, which lacks an assembly gap or repetitive structure. Notably, the region is inverted relative to Tak-2 (**D**).

Provisional gene IDs (Mp4g90480_P to Mp4g90960_P) were assigned to the duplicated genes.

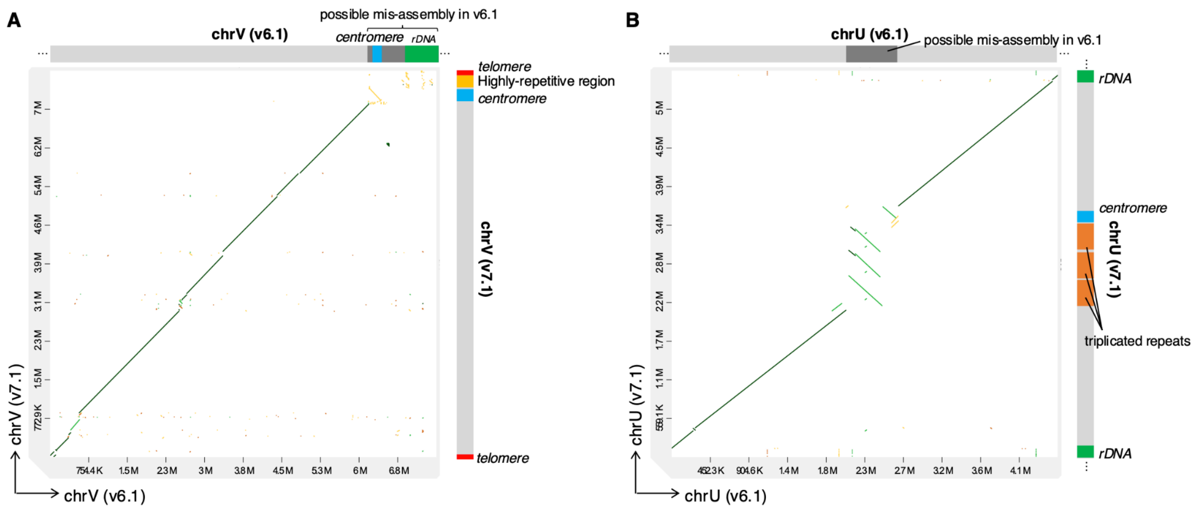


**Supplementary Fig. S7** Comparison of sex chromosomal sequences between ver. 7.1 (vertical axis) and ver. 6.1 (horizontal axis), visualized using D-Genies. Schematic diagrams of each sex chromosome from ver. 6.1 (top) and ver. 7.1 (right) are shown alongside the plots: **A**) Tak-1 chrV; **B**) Tak-2 chrU.

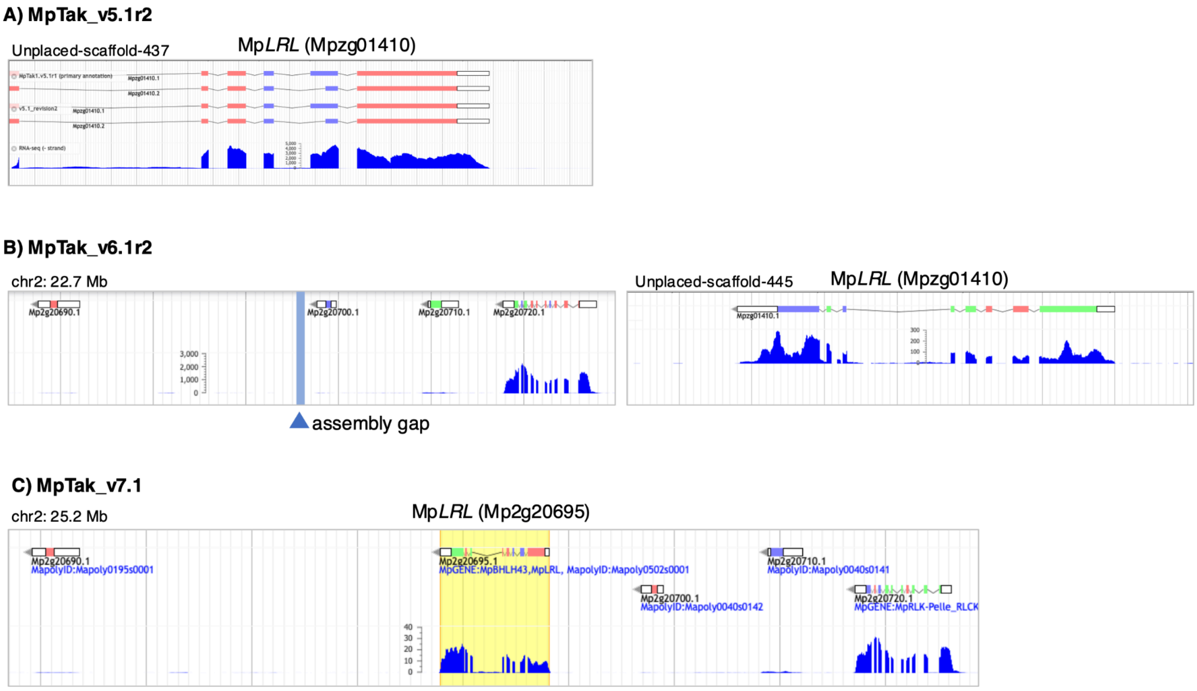


**Supplementary Fig. S8** Example of improved gene annotation using Mp*LRL* as a case study. **A**) In ver. 5.1 genome, Mp*LRL* is fragmented on unplaced scaffold 437 and assigned with Mpzg01410. **B**) In ver 6.1, The full gene sequence is recovered, but Mp*LRL* remains on unplaced scaffold 445. The corresponding region where Mp*LRL* is encoded in ver. 7.1 contains an assembly gap. **C**) In ver. 7.1, Mp*LRL* is located between Mp2g20690 and Mp2g20700 on chromosome 2, within a region that corresponds to an assembly gap in ver. 6.1, and is re-assigned as Mp2g20695.
